# SIZ1-mediated SUMOylation of ROS1 Enhances Its Stability and Positively Regulates Active DNA Demethylation in *Arabidopsis*

**DOI:** 10.1101/2020.03.05.978999

**Authors:** Xiangfeng Kong, Yechun Hong, Yi-Feng Hsu, Huan Huang, Xue Liu, Zhe Song, Jian-Kang Zhu

**Author notes:** Correspondence: Jian-Kang Zhu. These authors contributed equally to this work (Xiangfeng Kong, Yechun Hong).

## Abstract

The 5-methylcytosine DNA glycosylase/lyase REPRESSOR OF SILENCING 1 (ROS1)-mediated active DNA demethylation is critical for shaping the genomic DNA methylation landscape in *Arabidopsis*. Whether and how the stability of ROS1 may be regulated by post-translational modifications is unknown. Using a methylation-sensitive PCR (CHOP-PCR)-based forward genetic screen for *Arabidopsis* DNA hypermethylation mutants, we identified the SUMO E3 ligase SIZ1 as a critical regulator of active DNA demethylation. Dysfunction of SIZ1 leads to hyper-methylation at approximately one thousand genomic regions. SIZ1 physically interacts with ROS1 and mediates the SUMOylation of ROS1. The SUMOylation of ROS1 is reduced in *siz1* mutant plants. Compared to that in wild type plants, the protein level of ROS1 is significantly decreased, even though there is an increased level of *ROS1* transcripts in *siz1* mutant plants. Our results suggest that SIZ1 positively regulates active DNA demethylation by promoting the stability of ROS1 protein through SUMOylation.

**Short Summary:** The 5-methylcytosine DNA glycosylase/lyase REPRESSOR OF SILENCING 1 (ROS1) is indispensable for proper DNA methylation landscape in *Arabidopsis*. Whether and how the stability of ROS1 may be regulated by post-translational modifications is unknown. Here, we show that SIZ1-mediated SUMOylation of ROS1 enhances its stability and positively regulates active DNA demethylation.

## Introduction

As an important and conserved epigenetic mark, 5-methylcytosine DNA methylation takes part in various biological processes in plants and animals (Chang et al., 2020; Law and Jacobsen, 2010; Liu and Lang, 2020; Scott and Spielman, 2004; Zhang et al., 2018). In plants, DNA methylation occurs in all sequence contexts (CG, CHG, and CHH, H represents for A, T and G) which are established *de novo* via the RNA-directed DNA methylation (RdDM) pathway and maintained by specific mechanisms according to the sequence context. MET1 and CMT3 are responsible for maintaining DNA methylation at symmetric CG and CHG contexts, respectively (Law and Jacobsen, 2010). The CHH methylation is maintained by DRM2 through the RdDM pathway and by CMT2 (Zemach *et al*., 2013). DNA methylation is reversible and is determined by both methylation and demethylation processes (Zhu, 2009). The *Arabidopsis* 5-methylcytosine DNA glycosylase/lyase ROS1 is critical for pruning DNA methylation to keep a proper DNA methylation pattern genome-wide (Gong *et al*., 2002; Qian *et al*., 2012). ROS1 is recruited to specific genomic loci by the cooperation of the Increased DNA Methylation (IDM) and SWR1 complex (Lang *et al*., 2015; Li *et al*., 2015; Li *et al*., 2012; Nie *et al*., 2019; Qian *et al*., 2012; Wang *et al*., 2015). MET18, a cytosolic iron-sulfur assembly (CIA) pathway component, was identified as an important regulator for the enzymatic activity of ROS1 (Duan *et al*., 2015; Wang *et al*., 2016). Little is known about whether and how ROS1 may be regulated by post-translational modifications.

Mechanistically similar to ubiquitination, SUMOylation occurs through an ATP-dependent enzyme cascade, including heterodimeric E1 activating enzymes (SAE1/SAE2), E2 conjugating enzyme (SCE1), E3 ligase and SUMO proteases SENPs/Ulps (Augustine and Vierstra, 2018; Mukhopadhyay and Dasso, 2007; Seeler and Dejean, 2003). The SUMO E3 ligase can facilitate SUMO conjugation by E2 to the substrates and increase the substrate specificity (Gareau and Lima, 2010; Johnson, 2004). In *Arabidopsis*, the SUMO E3 ligase SIZ1 contains five domains including the SAP domain for nuclei acid binding, PHD domain for selecting targets and recognizing H3K4me3, PINIT motif required for the E3 ligase activity, SP-RING zinc finger domain for the E3 ligase activity and localization of SIZ1, and SXS motif for SUMO interaction (Cheong et al., 2009; Garcia-Dominguez et al., 2008; Miura et al., 2007a; Miura et al., 2020). Dysfunction of SIZ1 is reported to affect abiotic and biotic stress responses, phosphate starvation responses, flowering time and photomorphogenesis (Lin *et al*., 2016; Lois *et al*., 2003; Mazur *et al*., 2019; Miura *et al*., 2007a; Miura *et al*., 2007b; Miura *et al*., 2009; Miura *et al*., 2005).

Here, we show that SIZ1 positively regulates active DNA demethylation mainly through a ROS1-dependent pathway. Mutation of *SIZ1* leads to a genome-wide hyper-methylation phenotype similar to that in *ros1* mutants. We found that SIZ1 directly interacts with ROS1 and facilitates the SUMO modification of ROS1. Dysfunction of SIZ1 causes defects in ROS1 SUMOylation and a reduction in ROS1 protein level. Our study reveals an important connection between SUMOylation and active DNA demethylation in plants.

## Results and Discussion

### SIZ1 positively regulates active DNA demethylation independently of SA

To identify candidate genes involved in active DNA demethylation, we screened for T-DNA insertion mutants of *Arabidopsis thaliana* by CHOP-PCR based on DNA hyper-methylation phenotype at the 3’ region of *At1g26400* as previously described (Qian *et al*., 2012). Two mutants bearing T-DNA insertion in the SUMO E3 ligase gene *SIZ1* (SALK_065397/*siz1-2* and SALK_034008/*siz1-3*) showed a hyper-methylation phenotype (Fig. 1A and S1A). Locus-specific bisulfite sequencing result of the 3’ region of *At1g26400* indicated that in *ros1-4, siz1-2* and *siz1-3*, the methylation levels increased in all sequence contexts (i.e. CG, CHG and CHH), although the increase in non-CG methylation is not as pronounced as in CG methylation (Fig. 1B). CHOP-PCR tests on another three ROS1 target loci (*At1g26390, At1g26410* and *At4g18650*) were performed and all were found hyper-methylated in *siz1* mutants (Fig. 1C). Moreover, methylation analysis of the *ros1-4siz1-2* double mutant revealed that the hyper-methylation phenotypes of *siz1-2* and *ros1-4* were not additive (Fig. 1B and 1D), suggesting that SIZ1 and ROS1 may function in the same genetic pathway for active DNA demethylation.

**Figure 1.**
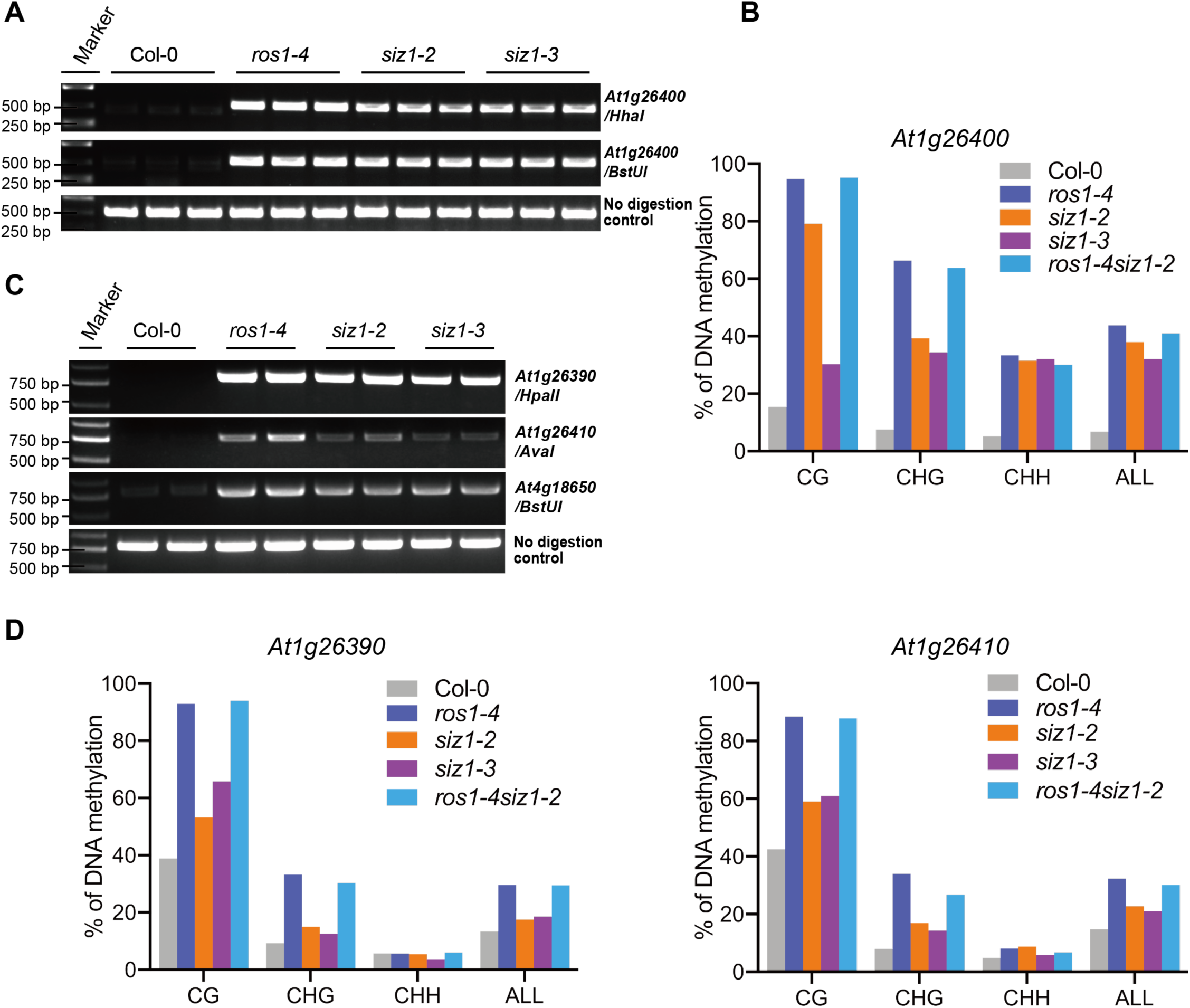
The *siz1* mutant plants show DNA hyper-methylation at multiple loci. (A) Analysis of DNA methylation status at the 3’ region of *At1g26400* by methylation-sensitive PCR (CHOP-PCR). Compared to that in wild type Col-0 plants, the methylation levels increased in *ros1-4, siz1-2* and *siz1-3* mutants. Undigested genomic DNA was used as a control. (B) Analysis of DNA methylation status at the 3’ region of *At1g26400* via locus-specific bisulfite sequencing. (C) Analysis of DNA methylation status at multiple loci by CHOP-PCR. Similar to those in *ros1-4* mutant, the methylation levels increased in *siz1-2* and *siz1-3* mutants. (D) Analysis of DNA methylation status at the indicated loci via locus-specific bisulfite sequencing. See also Figures S1 and S2.

Specific domains of SIZ1 control unique responses to different environmental stimuli (Cheong *et al*., 2009). To examine the function of the various domains of SIZ1 in regulating active DNA demethylation, we performed CHOP-PCR assay to analyze the SIZ1^WT^, SIZ1^sap^, SIZ1^phd^, SIZ1^pinit^, SIZ1^sp-ring^ and SIZ1^sxs^ plants and found that only expression of the wild type SIZ1 could rescue the hyper-methylation phenotype of *siz1-2* (Fig. S1B), demonstrating that intact SIZ1 protein is important for regulating DNA demethylation at the 3’ region of *At1g26400*. Dysfunction of SIZ1 leads to an obvious dwarf phenotype due to the elevated salicylic acid (SA) level (Lee *et al*., 2007). We introduced the *nahG* gene encoding a bacterial salicylate hydroxylase into *siz1-2* mutant plants to examine the effect of over-accumulated SA on DNA methylation. The expression of *nahG* rescued the dwarf phenotype of *siz1-2* but did not affect the DNA hyper-methylation phenotype (Fig. S2A and S2B), indicating that the hyper-methylation phenotype of *siz1-2* is not due to the elevated SA.

The indispensable role of SUMOylation in plant viability makes it difficult to analyze knock-out mutants of key genes in the SUMOylation pathway (Saracco *et al*., 2007). Over-expressing a mutated form of SCE1 containing the C94S substitution results in a dominant-negative effect, which impairs the SUMOylation process as previously reported (Tomanov *et al*., 2013). We generated SCE1(WT/C94S)-Flag over-expression plants (Fig. S3A), and found that plants over-expressing SCE1(C94S) showed a dwarf phenotype like *siz1* mutants (Fig. S3B). Only SCE1(C94S)-Flag over-expressing plants showed DNA hyper-methylation at the 3’ region of *At1g26400* similarly to *siz1-2* mutants (Fig. S3C), demonstrating that proper SUMOylation is important for active DNA demethylation.

### SIZ1 affects DNA methylation at over a thousand genomic regions

According to whole genome bisulfite sequencing data, 1,040 hyper-methylated DMRs (hyper-DMRs) and 183 hypo-methylated DMRs (hypo-DMRs) were identified in *siz1-2* mutant plants (Fig. 2A, Table S1 and Table S2). In accordance with locus-specific bisulfite sequencing results, whole genome bisulfite sequencing showed that *At1g26390, At1g26400, At1g26410* and *At4g18650* were all hyper-methylated compared to Col-0 (Fig. S4A), although only *At1g26410* was counted as a hyper-DMR by the stringent parameters used in this study. The hyper-DMRs in *siz1-2, ros1-4* and *rdd* (a triple mutant defective in *ROS1* and its paralogs *DML2* and *DML3*) mutants were distributed similarly in all five chromosomes (Fig. S4B and S4C). Analysis of hyper-DMRs in different genomic regions indicated that similar to that in *ros1-4*, ∼ 26% of hyper-DMRs in *siz1-2* were located in TE regions (Fig. S4D). Nine of the hyper-DMRs shared by *siz1* and *ros1* mutants were validated by CHOP-PCR (Fig. S5A and S5B).

**Figure 2.**
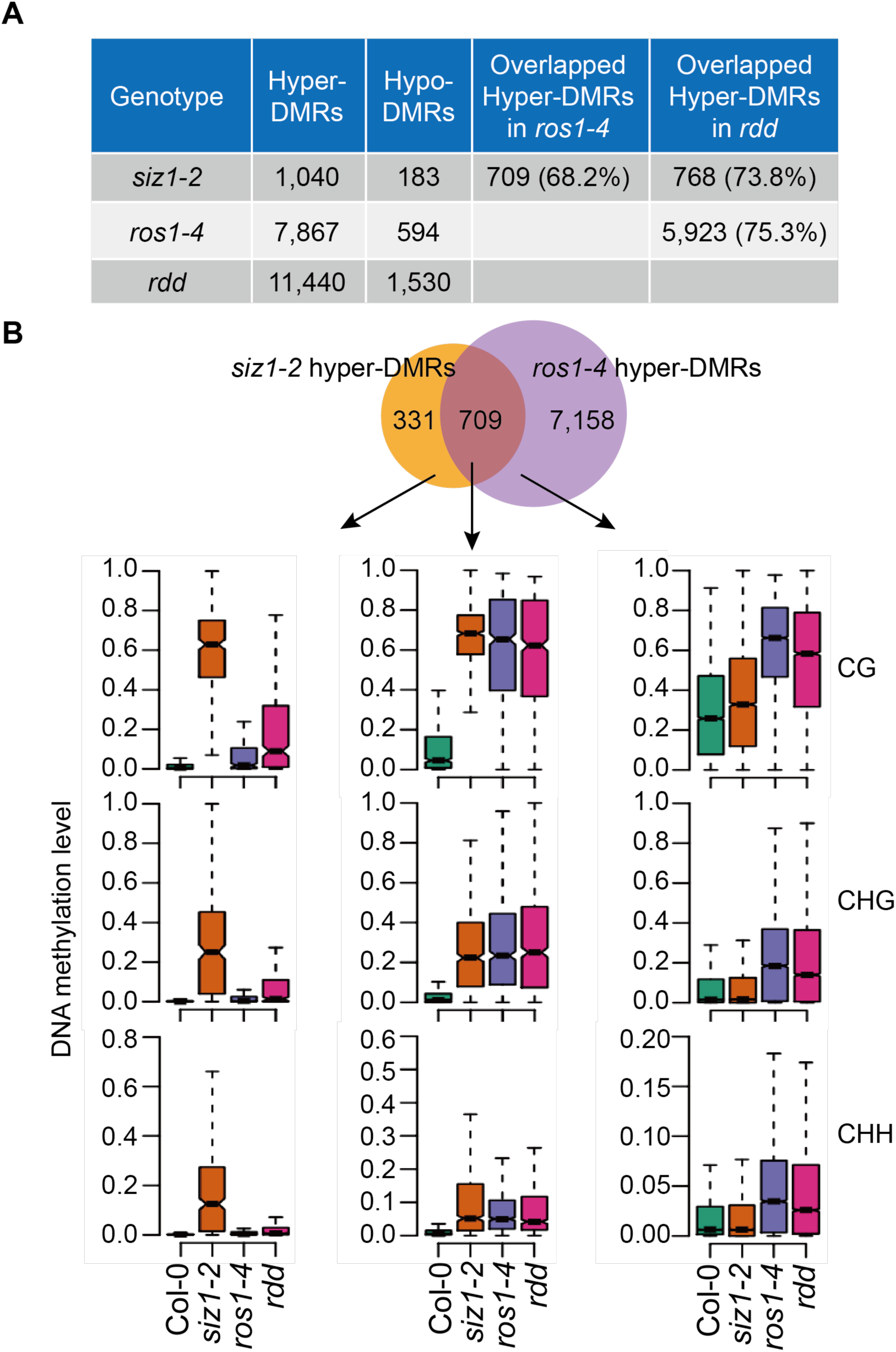
Effect of *SIZ1* mutation on genome-wide DNA methylation. (A) Numbers of hyper/hypo-DMRs identified in *siz1-2, ros1-4* and *rdd* mutants by whole-genome bisulfite sequencing, and the overlaps in hyper-DMRs between different mutants. (B) Venn diagram displaying the numbers of hyper-DMRs that are overlapping or unique in *siz1-2* and *ros1-4*. Box plots showing the distribution of average DNA methylation levels in CG, CHG and CHH contexts that were calculated from the indicated overlapping or unique hyper-DMRs. See also Figures S4, S5, S6 and Table S1-2.

Approximately 70% of the hyper-DMRs identified in *siz1-2* mutant plants overlapped with those in *ros1-4* and *rdd* mutant plants (Fig. 2A, 2B and Table S1). The DNA methylation level of the overlapping hyper-DMRs was increased in all cytosine contexts (Fig. 2B). However, the DNA methylation level of *ros1-4* specific hyper-DMRs was not increased in *siz1-2*, suggesting that SIZ1 affects a subset of ROS1 target loci. DNA methylation change is usually accompanied with perturbation of nearby genes’ expression (Harris et al., 2018; Liu and Lang, 2020; Yang et al., 2019; Zhao et al., 2019). To investigate whether the increased DNA methylation in *siz1-2* contributes to gene expression regulation, we analyzed 14 genes near the hyper-DMRs by RT-qPCR and found that six of the genes showed decreased transcript levels in *siz1* and *ros1* mutants (Fig. S6). These results suggested that SIZ1 may regulate the expression of these genes by affecting DNA methylation.

Among the 183 hypo-DMRs identified in *siz1-2* mutant plants, 179 (97.8%), 56 (30.6%), and 105 (57.4%) overlapped with hypo-DMRs in *met1, cmt3*, and *nrpd1*, respectively (Fig. S7 A-C and Table S2). For the overlapping hypo-DMRs of *siz1-2* and *met1*, their DNA methylation levels were decreased in all three cytosine contexts including CG, CHG and CHH (Fig. S7A). The DNA methylation levels of the overlapping hypo-DMRs between *siz1-2* and *cmt3* and between *siz1-2* and *nrpd1* were reduced mainly at CHG and CHH contexts, respectively (Fig. S7B and S7C). Even for the *siz1*-specific hypo-DMRs, the CHG and CHH DNA methylation levels were also decreased in *cmt3* and *nrpd1* (Fig. S7B and S7C). Heatmap analysis showed that the DNA methylation levels of most *siz1* hypo-DMRs were also reduced in *met1*, while the majority of non-CG methylation at *siz1* hypo-DMRs was decreased in *cmt3* and *nrpd1* (Fig. S7D). These results indicated that the DNA methylation at the *siz1-2* hypo-DMRs was mainly contributed by MET1, although CMT3 and RdDM may also contribute to the non-CG methylation in these regions. The potential contribution of CMT3 is in line with previous report of positive regulation of CMT3 activity by SUMOylation that is dependent on SIZ1 (Kim et al., 2015).

### SIZ1 physically interacts with ROS1

To examine if SIZ1 physically interacts with ROS1, yeast two-hybrid assays were performed. Only yeast cells carrying both SIZ1-AD and ROS1-BD grew well on the SD-Leu/-Trp/-His/-Ade media and turned blue with the supplement of X-IZ-Gal (Fig. 3A), indicating that SIZ1 interacts with ROS1 in yeast. The direct interaction between SIZ1 and ROS1 was confirmed by split-LUC and bimolecular fluorescence complementation (BiFC) assays in *Nicotiana benthamiana* leaves. In the split-LUC assay, strong LUC signals were detected only when cLUC-SIZ1 and ROS1-nLUC were co-expressed (Fig. 3B). The BiFC results showed a YFP fluorescence signal in the nucleus of *Nicotiana benthamiana* cells co-expressing SIZ1-cYFP and ROS1-nYFP (Fig. 3C), suggesting that SIZ1 interacts with ROS1 in the nucleus. Additionally, HA-SIZ1 could be co-immunoprecipitated with ROS1-Flag (Fig. 3D) when they were co-expressed in *Arabidopsis* protoplasts, which further confirmed the interaction between SIZ1 and ROS1. These results showed that SIZ1 physically interacts with ROS1 in the nucleus.

**Figure 3.**
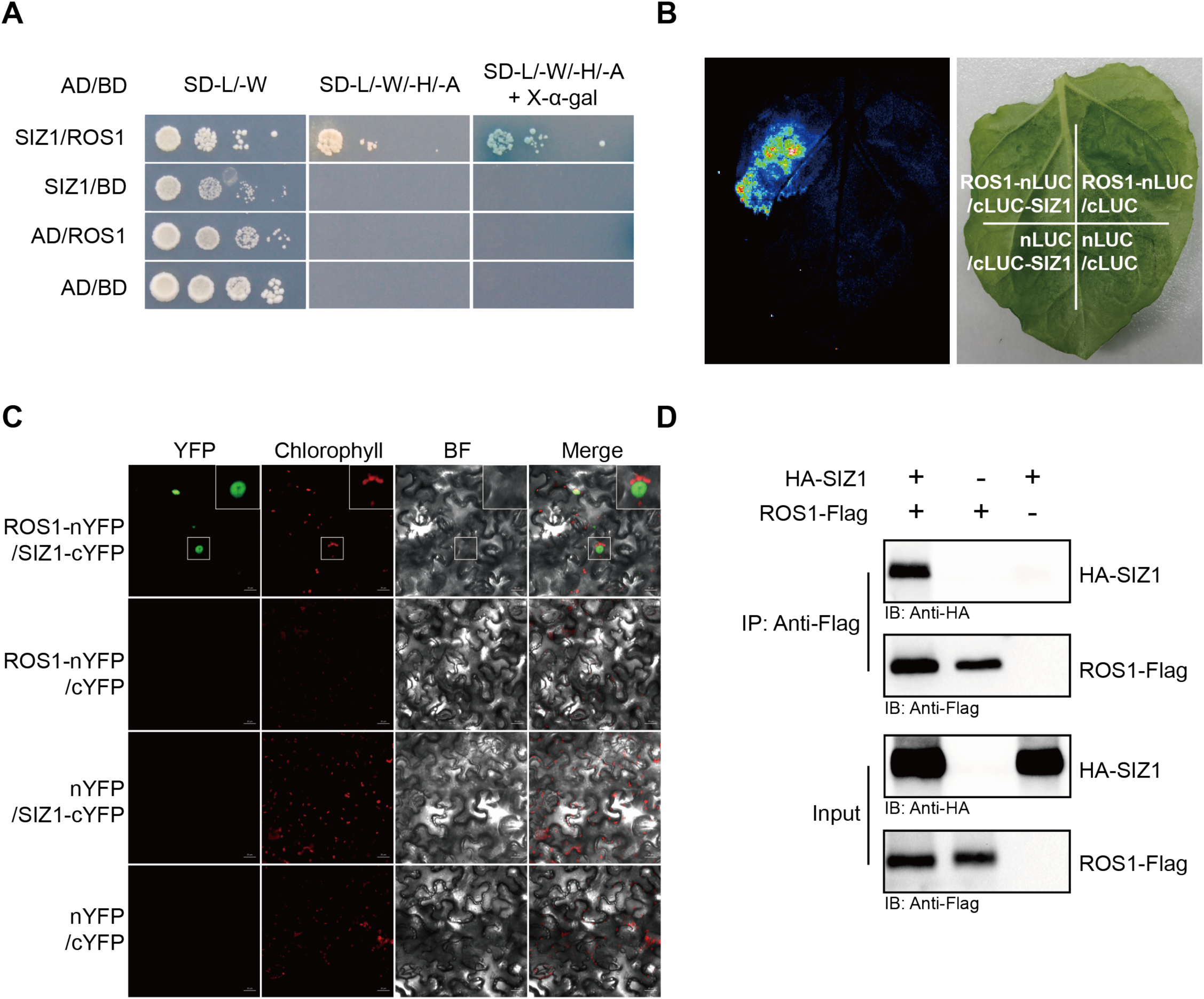
SIZ1 physically interacts with ROS1. (A) Yeast two-hybrid result showing an interaction between SIZ1 and ROS1. Full-length SIZ1 and ROS1 were fused to AD and BD, respectively. The different combination of recombinant plasmids and empty vectors were co-transformed into yeast cells. AD, Gal4 activation domain; BD, Gal4 binding domain. (B) Split-LUC assay showing an interaction between SIZ1 and ROS1 in *Nicotiana benthamiana* leaves. Firefly luciferase was fused with the coding sequences of *SIZ1* and *ROS1* at their N-terminus and C-terminus, respectively. (C) Bimolecular fluorescence complementation (BiFC) test indicating SIZ1 interaction with ROS1 in *Nicotiana benthamiana* leaves in the nucleus. Full-length SIZ1 and ROS1 were fused to YFP at their C-terminus. Chlorophyll, the autofluorescence of chlorophyll; BF, bright field; Scale bars, 20 µm. Inset, 4x magnification of boxed region. (D) Co-immunoprecipitation showing an association of SIZ1 with ROS1. HA-tagged SIZ1 and Flag-tagged ROS1 were expressed separately or co-expressed in *siz1-2* protoplasts. HA-SIZ1 and ROS1-Flag were detected by immuno-blotting using anti-HA and anti-Flag antibodies with crude lysate proteins and proteins immunoprecipitated via anti-Flag mAb-Magnetic beads.

### SIZ1 enhances the SUMOylation of ROS1 and stabilizes ROS1

The interaction between SIZ1 and ROS1 further prompted us to test whether ROS1 may be SUMOylated. Using the split-LUC assay, we found that only the combination of ROS1-nLUC or ROS1b-nLUC (510-1393 aa) with cLUC-SUMO1 showed strong LUC signals, indicating that ROS1 interacts with SUMO1 via ROS1 C-terminal region containing the DNA glycosylase and CTD domains (Fig. 4A). According to the *in vitro* SUMOylation assay described previously (Okada *et al*., 2009) and SUMOylation site prediction (Fig. S8), ROS1c (1-290 aa, containing K133) and ROS1d (794-1063 aa, containing K806/K812/K851/K1051) fused with the T7-tag were generated. Immunoblot analysis detected a shift of protein band to higher molecular weight when ROS1d but not ROS1c was co-expressed with SUMO E1, SUMO E2 and the mature SUMO1GG (Fig. 4B). A similar result was obtained when T7-ROS1c/ROS1d was replaced with GST-ROS1c/ROS1d-Myc in the assay (Fig. S9A and S9B), suggesting that ROS1 can be SUMOylated *in vitro*, and the SUMOylation was mainly in the DNA glycosylase domain.

**Figure 4.**
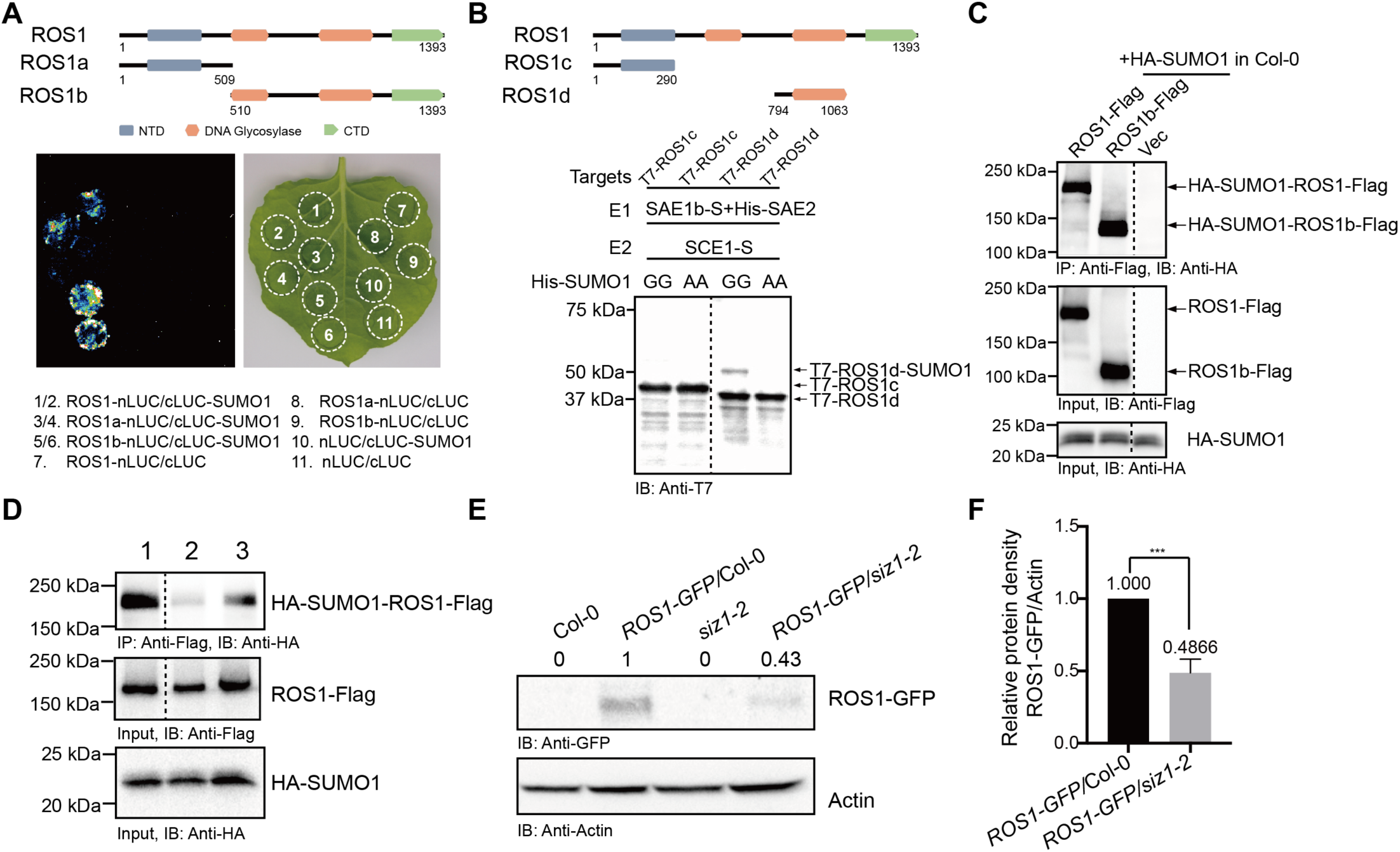
SIZ1 mediated SUMOylation of ROS1 enhances its stability. (A) The interaction of SUMO1 and ROS1 was tested using the split-LUC assay. The C-terminal end of firefly luciferase was fused to the N-terminus of SUMO1. ROS1 was fused to the N-terminal end of firefly luciferase. The schematic diagram shows the full-length ROS1 and truncated ROS1 (ROS1a: 1-509 aa, ROS1b: 510-1393 aa). NTD, N-terminal domain; CTD, C-terminal domain. (B) SUMOylation of ROS1 was tested using the *Arabidopsis* SUMOylation pathway reconstitution system in *E. coli*. Different combinations of the three plasmids encoding *Arabidopsis* SUMOylation machinery proteins and truncated ROS1 indicated above each lane were co-transformed into *E. coli* BL21(DE3). Immuno-blotting was used to detect truncated ROS1 and its SUMOylated forms using anti-T7 antibodies in crude lysate proteins. SUMOylated ROS1 shifting to a larger molecular weight band was observed only in the lane using the mature wild type form of SUMO1 (referred to as SUMO1GG), but not in lanes using mutated form of SUMO1 (referred to as SUMO1AA). The schematic diagram indicates the truncated ROS1 (ROS1c: 1-290 aa containing K133), ROS1d: 794-1063 aa containing K806, K812, K851 and K1051). (C) *In vivo* SUMOylation of ROS1. ROS1-Flag/ROS1b-Flag and HA-SUMO1 were co-expressed in Col-0 protoplasts. The SUMOylated ROS1 was detected via immuno-blotting using anti-HA antibodies in proteins immunoprecipitated by anti-Flag mAb-Magnetic beads. Expression of HA-SUMO1 with empty vector served as a negative control. (D) SUMOylation of ROS1 was reduced in *siz1-2*. ROS1-Flag (lane1) / ROS1(5K/Rs)-Flag (lane2) and HA-SUMO1 were co-expressed in Col-0 protoplasts. ROS1-Flag and HA-SUMO1 were co-expressed in *siz1-2* protoplasts (lane3). SUMOylation in mutant ROS1 with the 5 predicted SUMOylation sites mutated by lysine to arginine substitutions (K133R/K806R/K812R/K851R/K1051R) was reduced (lane2). SUMOylation of ROS1 was substantially decreased in *siz1-2* compared to that in Col-0 (lane 3). (E) Immunodetection of ROS1 by an anti-GFP antibodies in GFP knock-in lines with GFP integrated in the endogenous *ROS1* locus in Col-0 and *siz1-2* background. Col-0 and *siz1-2* without the GFP served as negative controls. Actin was used as a loading control. The protein level of ROS1 relative to Actin shown above the lanes was quantified using Image Lab software (version 5.2.1, Bio-Rad). (F) Statistical analysis of the ROS1 protein levels in *siz1-2* mutant (n=3). ^***^ *P*< 0.001, determined by Student’s *t*-test. See also Figures S8, S9, S10 and S11.

An *in vivo* SUMOylation assay was performed to confirm the SUMOylation of ROS1. A shift of protein band to higher molecular weight was detected when ROS1-Flag or ROS1b-Flag was co-expressed with HA-SUMO1 in Col-0 protoplasts. The higher molecular weight band was confirmed to contain both ROS1 and SUMO1 by LC-MS/MS analysis (Fig. 4C and S10). The band corresponding to SUMO1-modified ROS1 was substantially weaker when the proteins were expressed in *siz1-2* protoplasts (Fig. 4D), indicating that SIZ1 enhances the SUMO1 modification of ROS1. The lysine to arginine substitution can block the SUMOylation on substrate proteins (Gostissa *et al*., 1999), so we mutated 5 predicted SUMOylation sites to generate ROS1(5K/Rs)-Flag and co-expressed the mutant protein with HA-SUMO1 in Col-0 protoplasts. The SUMO1 modification of ROS1 was blocked by these 5 lysine to arginine substitutions (Fig. 4D), demonstrating that ROS1 is SUMOylated at one or more of these 5 predicted lysine residues.

Finally, we examined the effect of SIZ1-mediated SUMOylation on ROS1 by analyzing the protein level of ROS1 in *siz1-2* mutant. An *ROS1-GFP*/*siz1-2* line was obtained by crossing *siz1-2* to the *GFP* knock-in line *ROS1-GFP*/Col-0 generated in a previous study (Miki *et al*., 2018). Although *ROS1* transcript level significantly increased in *siz1-2* mutant (Fig. S11A and S11B), the immunoblotting result showed that the ROS1-GFP protein was substantially deceased in *siz1-2* mutant plants (Fig. 4E and 4F). Together, these results support that SIZ1 enhances the stability of ROS1 by facilitating the SUMOylation of ROS1.

The role of SUMO modification in active DNA demethylation has been reported in mammals (McLaughlin *et al*., 2016; Steinacher *et al*., 2019; Waters *et al*., 1999). In *Arabidopsis*, some components of the RdDM pathway and base excision repair (BER) pathway were found to be SUMOylated or interact with SUMOs (Elrouby *et al*., 2013; Kim *et al*., 2015; Miller *et al*., 2010; Rytz *et al*., 2018). In this study, we showed that the SUMO E3 ligase SIZ1 positively regulates active DNA demethylation by mediating SUMOylation of ROS1, leading to enhanced ROS1 stability (Fig. 4A-4D, S8-S10). Additionally, SIZ1 may regulate DNA methylation by facilitating the SUMOylation of DNA methyltransferases such as CMT3 and other methylation regulators, which are possibly responsible for the hypo-DMRs in *siz1* mutant plants (Fig. 2A and S7) (Augustine and Vierstra, 2018; Kim et al., 2015).

In mammals, the DNA glycosylase TDG is SUMOylated to enhance its enzymatic turnover, leading to more efficient active DNA demethylation (McLaughlin *et al*., 2016; Waters *et al*., 1999). Here, we showed that the important DNA glycosylase ROS1 in *Arabidopsis* can also be SUMOylated and this is facilitated by SUMO E3 ligase SIZ1 (Fig. 4A-4D, S8-S10). The accumulation of ROS1 protein is reduced while *ROS1* transcripts are increased in *siz1* mutant plants (Fig. 4E, 4F and S11), which may be due to feedback regulation through the DNA methylation monitoring sequence at the *ROS1* gene promoter as reported in previous studies (Lei et al., 2015; Qian et al., 2014; Xiao et al., 2019; Zheng et al., 2008). Only a certain subset of ROS1 target loci was affected in *siz1* mutant plants, which may be caused by the distinct requirements of ROS1 function at different genomic regions. Since the possible SUMOylation sites are located in the DNA glycosylase domain that is critical for 5-methylcytosine excision activity of the 5-methylcytosine DNA glycosylases (Gehring et al., 2006; Mok et al., 2010; Ortega-Galisteo et al., 2008), the role of SUMOylation on the enzymatic activity or turnover of ROS1 needs to be tested once the precise SUMOylated residues are determined. In addition, it will be interesting to study the role of SIZ1-mediated ROS1 SUMOylation in regulating ABA responses, considering the similar DNA hyper-methylation in the promoter region of *DOLG4* displayed by both the *siz1* and *ros1-4* mutants (At4g18650, Fig. 1C and S4A) (Zhu et al., 2018). Since ROS1 has been suggested to be modified by ubiquitination, whether SUMOylation of ROS1 competitively affects the ubiquitination of ROS1 to reduce its degradation remains to be determined in the future (Hay, 2005; Maor et al., 2007; Miura et al., 2007b; Ulrich, 2005).

## Materials and Methods

### Plant materials and CHOP-PCR-based screening

The *Arabidopsis siz1-2* (SALK_065397) and *siz1-3* (SALK_034008) mutant lines were obtained from the Arabidopsis Biological Resource Center (ABRC, http://www.arabidopsis.org). The coding sequence of *SCE1*(WT) or *SCE1*(C94S) were cloned into pCAMBIA1305 vector and introduced into Col-0 plants to generate SCE1 (WT) and SCE1 (C94S) over-expression lines. Seedlings were grown on 1/2 Murashige and Skoog (MS) agar plates at 22 **°**C under 16 h light/8 h dark photoperiod. Plants including *Arabidopsis* and *Nicotiana benthamiana* were grown in soil at 22 **°**C under a 16 h light/8 h dark photoperiod. Screening for mutants was as described (Qian *et al*., 2012). Primers are listed in Table S3.

### Locus-specific bisulfite sequencing

Experiment was performed as described (Li et al., 2020). Primers used for PCR were listed in Table S3.

### Whole genome bisulfite sequencing and data analysis

Fourteen-day-old seedlings were used for genomic DNA extraction using DNeasy Plant Kit (Qiagen) following the manufacturer’s protocols. Bisulfite treatment, library construction, and deep sequencing were performed by the Genomics Core Facility at the Shanghai Center for Plant Stress Biology, China.

Adaptor and low-quality sequences (q < 20) were trimmed to generate clean reads that were then mapped to the TAIR 10 genome using BSMAP (Bisulfite Sequence Mapping Program) and allowing two mismatches. DMRs were counted according to Qian et al. (2012) with some modifications. Briefly, the cytosines with >= 4X coverage were considered. DMCs were identified if the *P*-value calculated by two-tailed Fisher’s Exact test was < 0.01. Genome was divided into 1kb regions, in which the number of DMCs was counted. A region with at least 5 DMCs was considered as an anchor region, of which the actual boundary was adjusted as the locations of the first DMC and last DMC. The anchor regions were combined into a larger region if the distance between two anchor regions was <= 1kb. A DMR was reported if the larger region contained at least 7 DMCs.

### Yeast two-hybrid assay

Yeast two-hybrid assay was carried out as described previously (Hong *et al*., 2020). Briefly, the coding sequence of *ROS1* and *SIZ1* were cloned into pGBKT7 and pGADT7 (Clontech), respectively. Different combinations of bait and prey plasmids were co-transformed as indicated.

### Split-LUC assay and Bimolecular fluorescence complementation (BiFC) assay

The split-LUC assay and BiFC assay were performed as described (Hong *et al*., 2020). Briefly, the full-length coding sequences of *ROS1, SIZ1* and *SUMO1* were amplified to generate cLUC-SIZ1, cLUC-SUMO1, ROS1-nLUC, SIZ1-p2YC and ROS1-p2YN constructs. The truncated ROS1a/ROS1b-nLUC constructs were generated from ROS1-nLUC using indicated primers (Table S3).

### Co-Immunoprecipitation assay

The full-length coding sequences of *ROS1* and *SIZ1* were cloned into pUC18-Flag and pUC18-HA vectors separately, then co-expressed in Col-0 protoplasts by polyethylene glycol (PEG400)-mediated transformation as previously described (Yoo *et al*., 2007). The protoplasts were collected and lysed after an incubation for 20 h at 22 °C. Fifty μl of anti-Flag mAb-Magnetic beads (MBL International) were added to the crude lysate supernatant and gently rotated for 2 h at 4 °C. The immunoprecipitated proteins were detected by Western-blotting using anti-HA antibody (Roche) and anti-Flag antibody (Sigma).

### SUMOylation assay and mass spectrometry

The *in vitro* SUMOylation assay was performed as previously described (Okada *et al*., 2009). The pET28a_ ROS1c / ROS1d expressing N-terminal T7-tagged variant ROS1 (ROS1c: 1-290 aa, ROS1d: 794-1063 aa) and pGEX4T1_ROS1c / ROS1d expressing GST-ROS1c/ROS1d-Myc were generated. The SUMOylation was analyzed by Western-blotting using anti-T7 antibody (Abcam) or anti-Myc antibody (Millipore).

*In vivo* SUMOylation assay was conducted as described (Zheng *et al*., 2012).The coding sequences of *ROS1* or *SUMO1* were cloned into pUC18-Flag or pUC18-HA respectively and co-expressed in Col-0 or *siz1-2* protoplasts. The protoplasts were collected after 20 h of incubation at 22 °C. The proteins were immunoprecipitated by anti-Flag mAb-Magnetic beads (MBL International) detected by Western-blotting using anti-HA antibody (Roche) and anti-Flag antibody (Sigma).

For LC-MS/MS analysis, the anti-Flag mAb-Magnetic beads immunoprecipitated proteins were separated in SDS-PAGE gel followed by silver staining using Fast Silver Stain Kit (Beyotime). The bands of ROS1 and SUMOylated ROS1 were cut into pieces according to their molecular weights and used for LC-MS/MS analysis at BGI.

### RNA extraction and Real-Time quantitative RT-PCR

RNA extraction and gene transcripts level determination by RT-qPCR were performed using the ChamQ SYBR qPCR Master Mix (Vazyme Biotech Co.,Ltd) according to (Hong *et al*., 2020). *UBQ10* and *ACT2* were used as internal control. The primers used were listed in Table S3.

### Accession numbers

Sequence data from this article can be found in The Arabidopsis Information Resource (http://www.arabidopsis.org/) under the following accession numbers: *SIZ1* (At5g60410), *ROS1* (At2g36490), *SUMO1* (At4g26840), *SCE1* (At3g57870), *TUB8* (At5g23860), *UBQ10* (At4g05320), *ACT2* (At3g18780). The methylome data used in this study can be found in NCBI GEO by the following accession numbers: GSE152425 (*siz1-2*), GSE33071 (Qian et al., 2012) (Col-0, *ros1-4* and *rdd*), GSE39901 (Stroud et al., 2013) (*met1, cmt3* and *nrpd1*).

## Supplemental Items

**Supplemental Figure1**. Characterization of *siz1* mutants.

**Supplemental Figure2**. Hyper-methylation caused by *siz1* mutation is not due to the elevated SA.

**Supplemental Figure 3**. Plants over-expressing dominant-negative SUMO E2 SCE1(C94S) show DNA hypermethylation at *At1g26400* loci.

**Supplemental Figure 4**. Analysis of DNA methylation pattern of *siz1-2* by whole genome bisulfite sequencing.

**Supplemental Figure 5**. Examples of hyper-methylated regions.

**Supplemental Figure 6**. DNA hypermethylation represses the expression of nearby genes in *siz1* mutants.

**Supplemental Figure 7**. Features of *siz1-2* hypo-DMRs.

**Supplemental Figure 8**. Predicted SUMOylation sites in ROS1.

**Supplemental Figure 9**. ROS1 is SUMOylated.

**Supplemental Figure 10**. Detection of SUMOylated ROS1 by LC-MS/MS.

**Supplemental Figure 11**. *ROS1* transcript level in the *siz1* mutants.

**Supplemental Table 1**. List of hyper-DMRs identified in *siz1-2, ros1-4* and *rdd* mutants.

**Supplemental Table 2**. List of hypo-DMRs of *siz1-2, met1, cmt3* and *nrpd1* mutants.

**Supplemental Table 3**. Primers used in this study.

## Author contributions

J.-K.Z., X.K. and Y.-F.H conceived the research. X.K., Y.H., Y.-F.H., X.L. and Z.S. performed experiments. H.H. analyzed the methylome data. X.K., Y.H., Y.-F.H. and J.-K.Z. wrote the manuscript.

## Acknowledgements

We thank Dr. Katsunori Tanaka for kindly providing the constructs for SUMOylation test in *E*.*coli* (pCDFDuet-AtSUMO1(AA/GG)-AtSCE1, pACYCDuet-AtSAE1b-AtSAE2), Dr. Dae-Jin Yun for kindly providing the transgenic plants of *SIZ1* carrying point mutations at specific domains (SIZ1^sap^, SIZ1^phd^, SIZ1^pinit^, SIZ1^sp-ring^, SIZ1^sxs^), Drs. Chao-Feng Huang and Jie Zhang for their technical support and suggestions. This work was supported by the Chinese Academy of Sciences to J.-K.Z.

