## Supplemental Information for "SIZ1-mediated SUMOylation of ROS1 Enhances Its Stability and Positively Regulates Active DNA Demethylation in *Arabidopsis*"

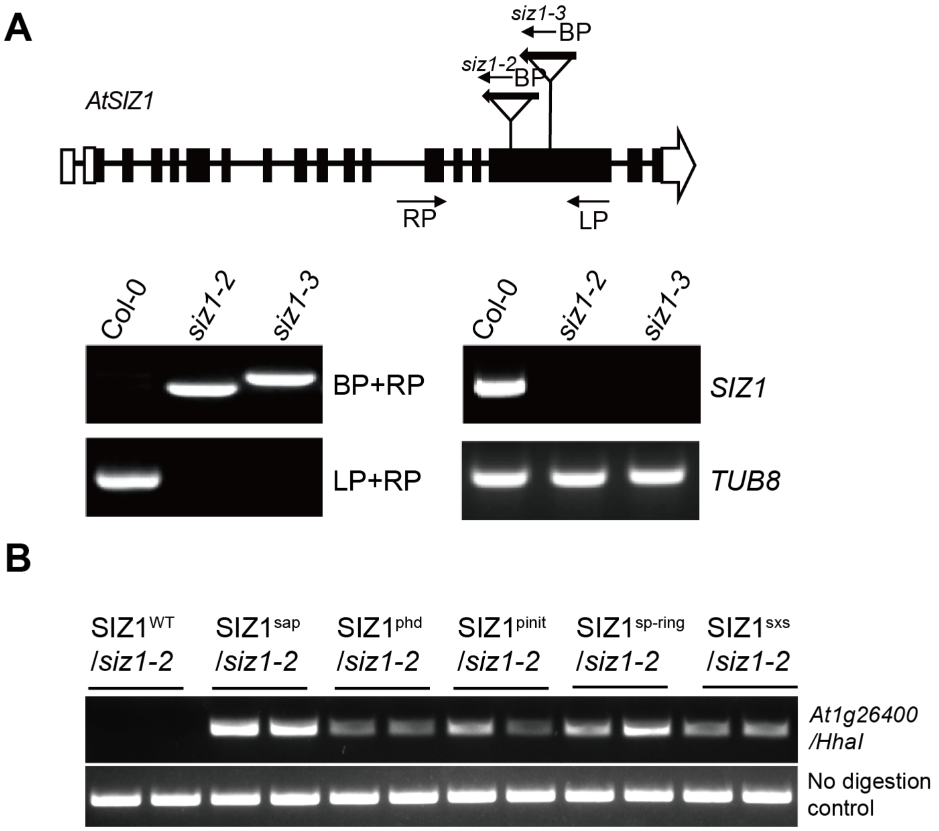


**Supplemental Figure 1. Characterization of *siz1* mutants.** (A) Schematic diagram showing the T-DNA insertion position at *SIZ1* locus. Black rectangles and white rectangles represent exons and UTRs, respectively. Genotyping and RT-PCR results showing both the two *siz1* mutants are homozygous and knock-out allele. (B) DNA methylation level analysis of indicated *siz1-2* transformants. The hyper-methylation phenotype caused by *siz1-2* mutation was recovered only in wild-type *SIZ1* transgenic plants. CHOP-PCR results for analysis of methylation status at *At1g26400* loci in indicated mutants and transgenic plants were shown. The no digestion was served as a loading control.

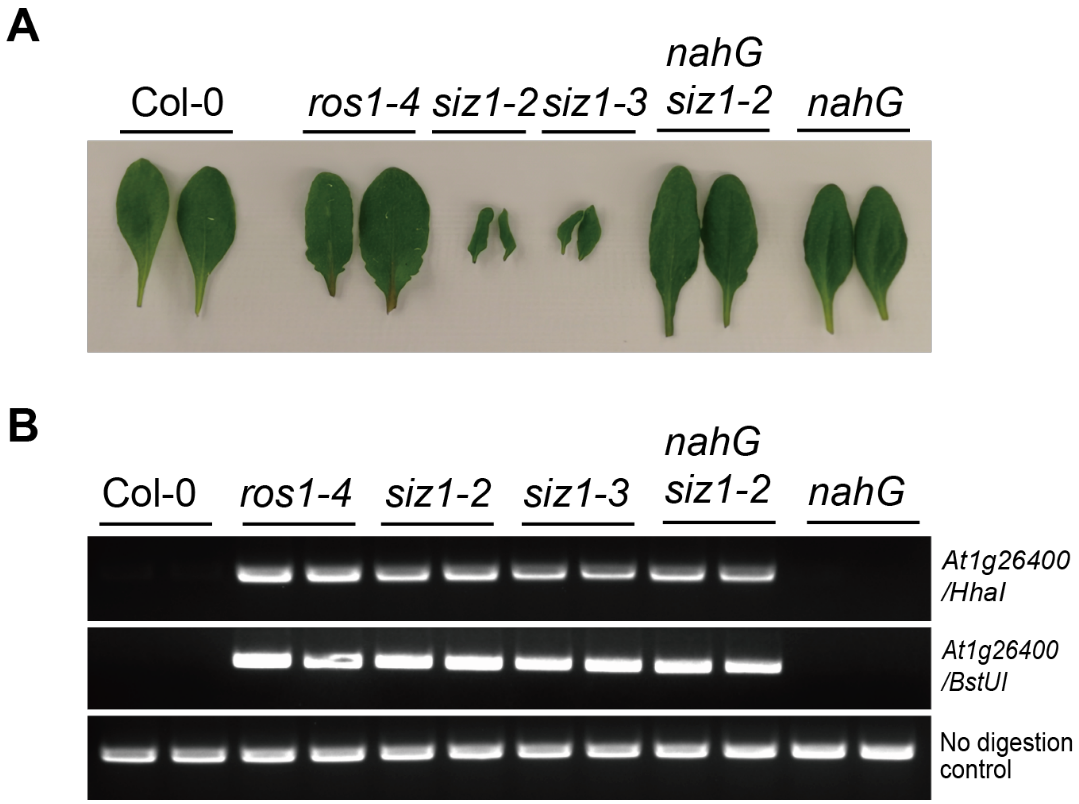


**Supplemental Figure 2. Hyper-methylation caused by *siz1* mutation is not due to the elevated SA.** (A) The dwarf phenotype of *siz1* mutants was rescued by introduction of *nahG*. (B) The hyper-methylation phenotype caused by *siz1* mutation could not be rescued by introducing *nahG* into *siz1-2*. CHOP-PCR results for analysis of methylation status at the 3' region of *At1g26400* in indicated mutants and transgenic plants were shown. The no digestion was served as a loading control.

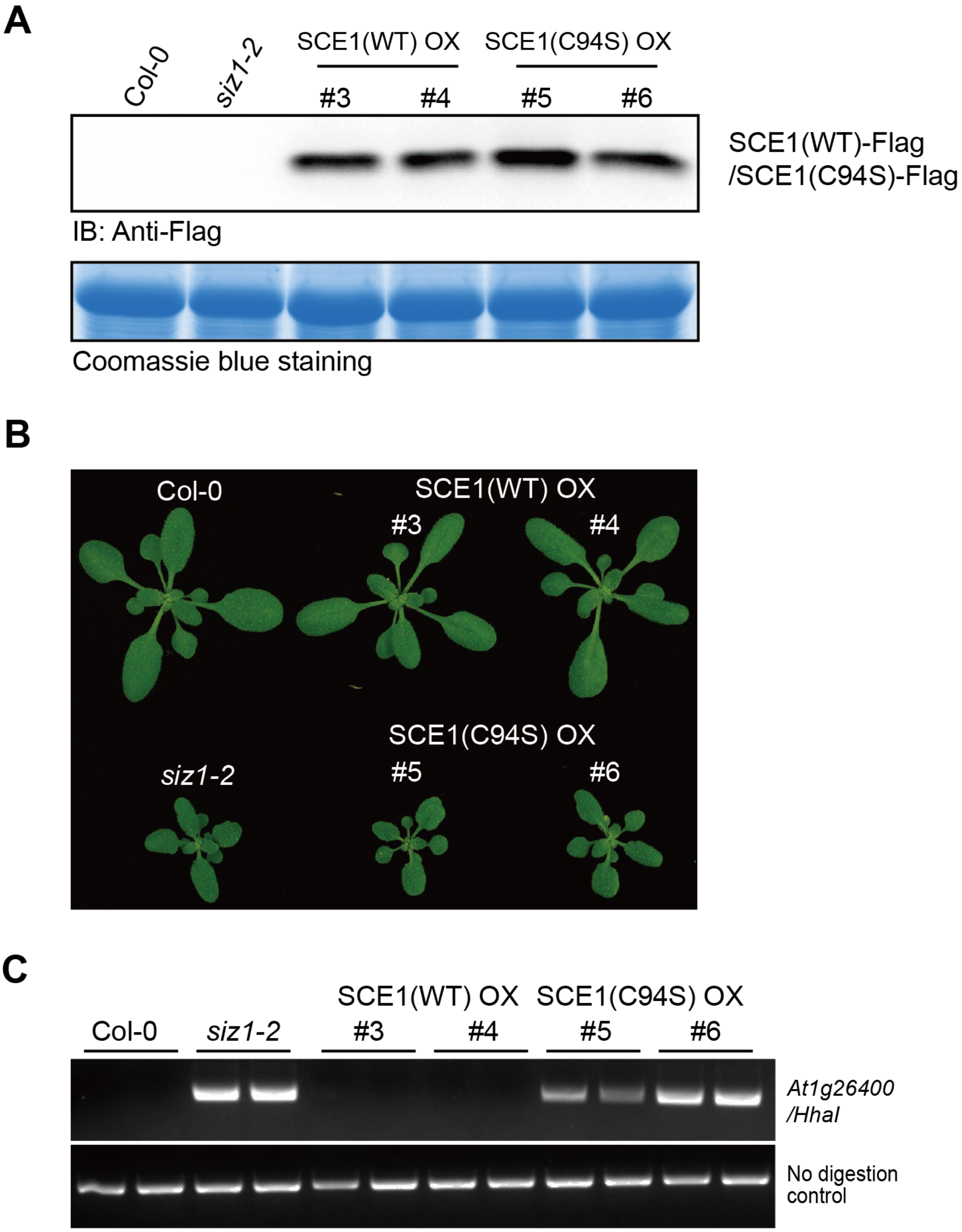


**Supplemental Figure 3. Plants over-expressing dominant-negative SUMO E2 SCE1(C94S) show DNA hypermethylation at *At1g26400* loci.** (A) Immunodetection of SCE1(WT)-Flag/SCE1(C94S)-Flag. SCE1(WT)-Flag/SCE1(C94S)-Flag was detected by anti-Flag antibody with crude lysate from transgenic plants. Col-0 and *siz1-2* were served as negative control. (B) Growth phenotype of indicated genotype plants. (C) Over-expression of SCE1(C94S) in Col-0 plants led to hypermethylation at *At1g26400* loci similar to that in *siz1* mutants. CHOP-PCR results for analysis of methylation status at *At1g26400* loci in indicated mutants and transgenic plants were shown. The no digestion was served as a loading control. OX, over-expression.

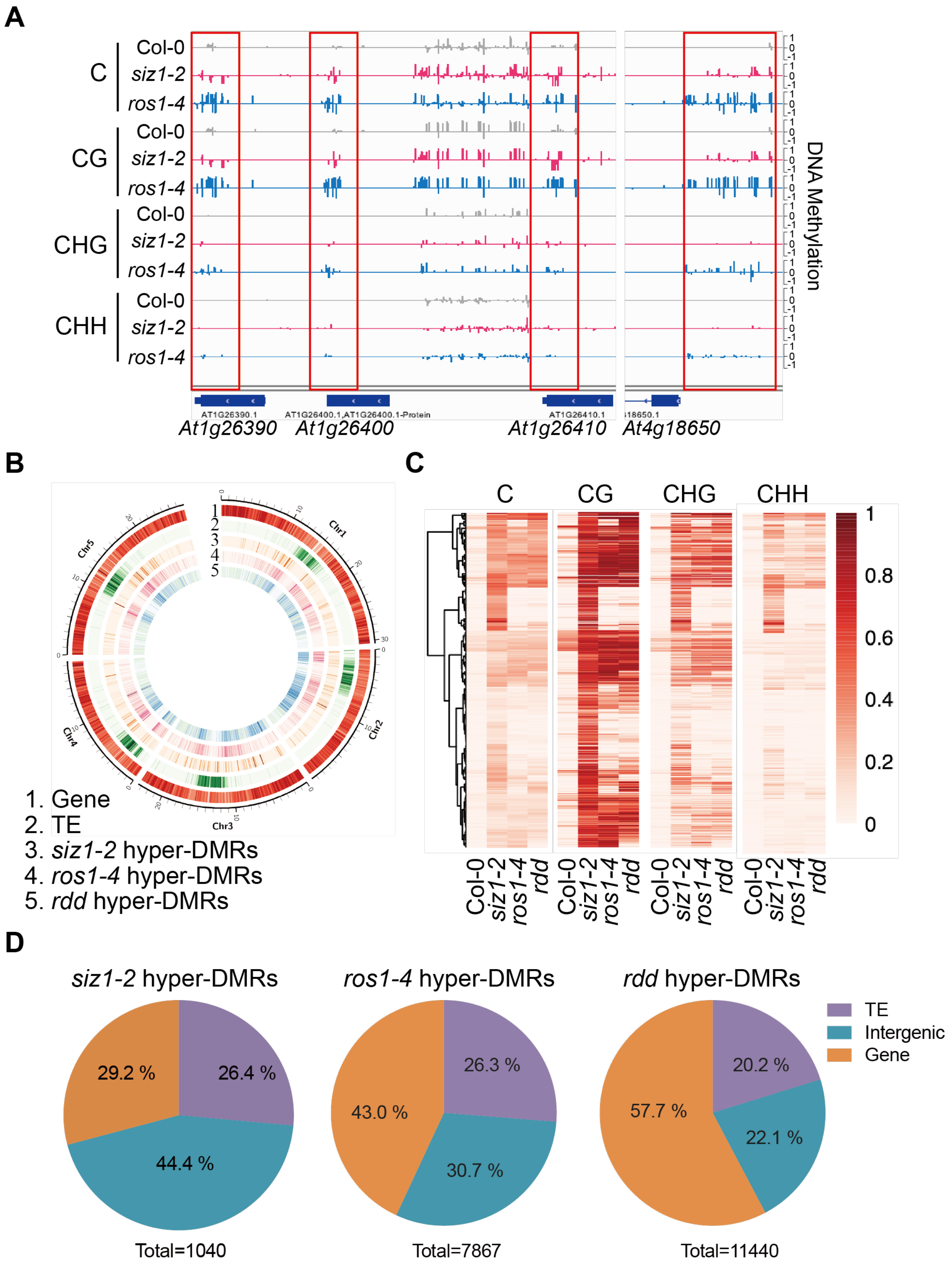


**Supplemental Figure 4. Analysis of hyper-methylation pattern of *siz1-2* by whole genome bisulfite sequencing.** (A) Integrated Genome Viewer (IGV) snapshot showing the genome loci and DNA methylation level of CHOP-PCR markers (*At1g26390*, *At1g26400*, *At1g26410* and *At4g18650*) in Col-0, *siz1-2* and *ros1-4*. Hyper-methylated regions were highlighted by red-box. (B) Density plots of DNA methylation showing genomic distribution of hyper-DMRs in *siz1-2*, *ros1-4* and *rdd* mutants. Density of gene and transposable element (TE) on five chromosomes were indicated. (C) Heatmap showing the DNA methylation level at different cytosine contexts in *ros1-4* and *rdd* within those regions that are hyper-methylated in *siz1-2*. (D) Composition of hyper-DMRs in *siz1-2*, *ros1-4* and *rdd* mutants.

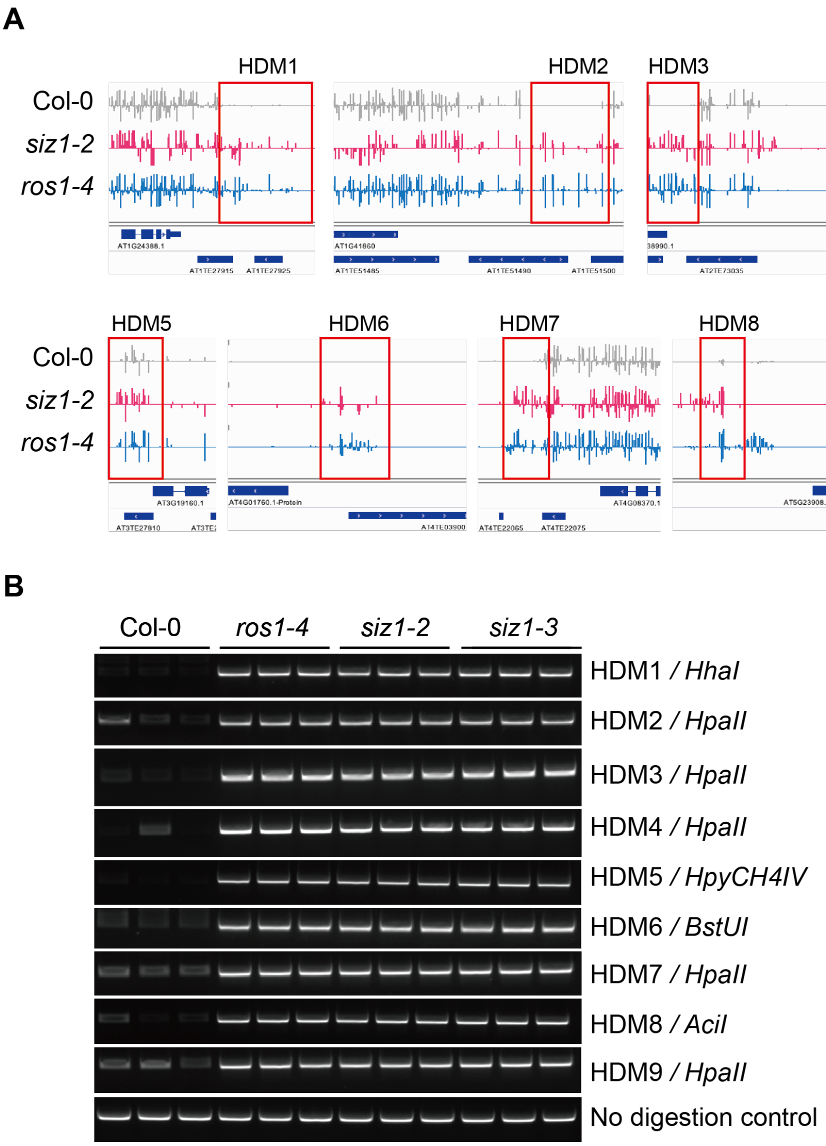


**Supplemental Figure 5**. **Examples of hyper-methylated regions.** (A) IGV snapshot showing the genome loci and DNA methylation level at different loci in indicated genotypes. HDM, short for Hyper-DNA Methylation. (B) Validation of whole genome bisulfite sequencing results by CHOP-PCR.

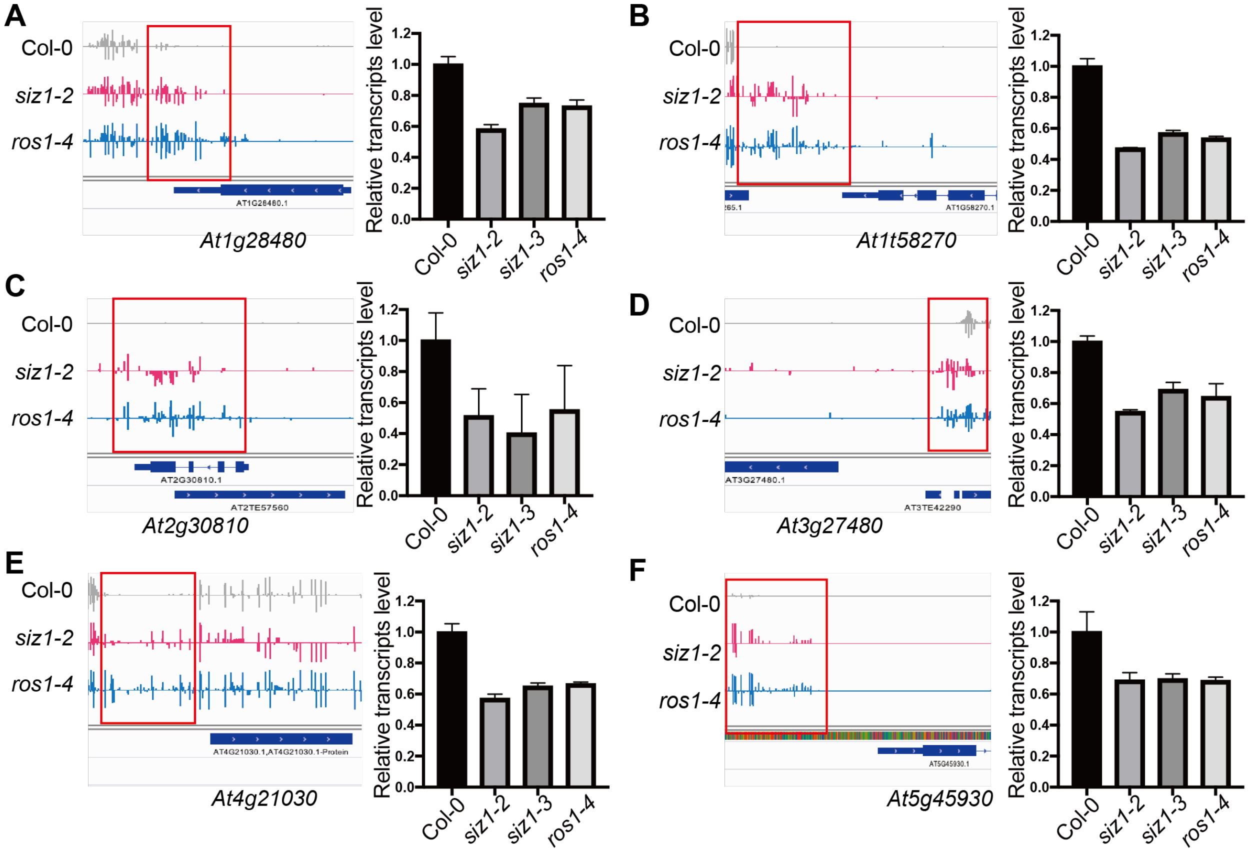


**Supplemental Figure 6. DNA hypermethylation represses the expression of nearby genes in *siz1* mutants.** (A-F) Left panel: IGV snapshot showing the DNA methylation level of hyper-DMRs in *siz1-2* and *ros1-4*. Hyper-methylated regions were highlighted by red-box. Right panel: The relative transcripts level of nearby genes. The error bars indicates the mean ± SD (n=3).

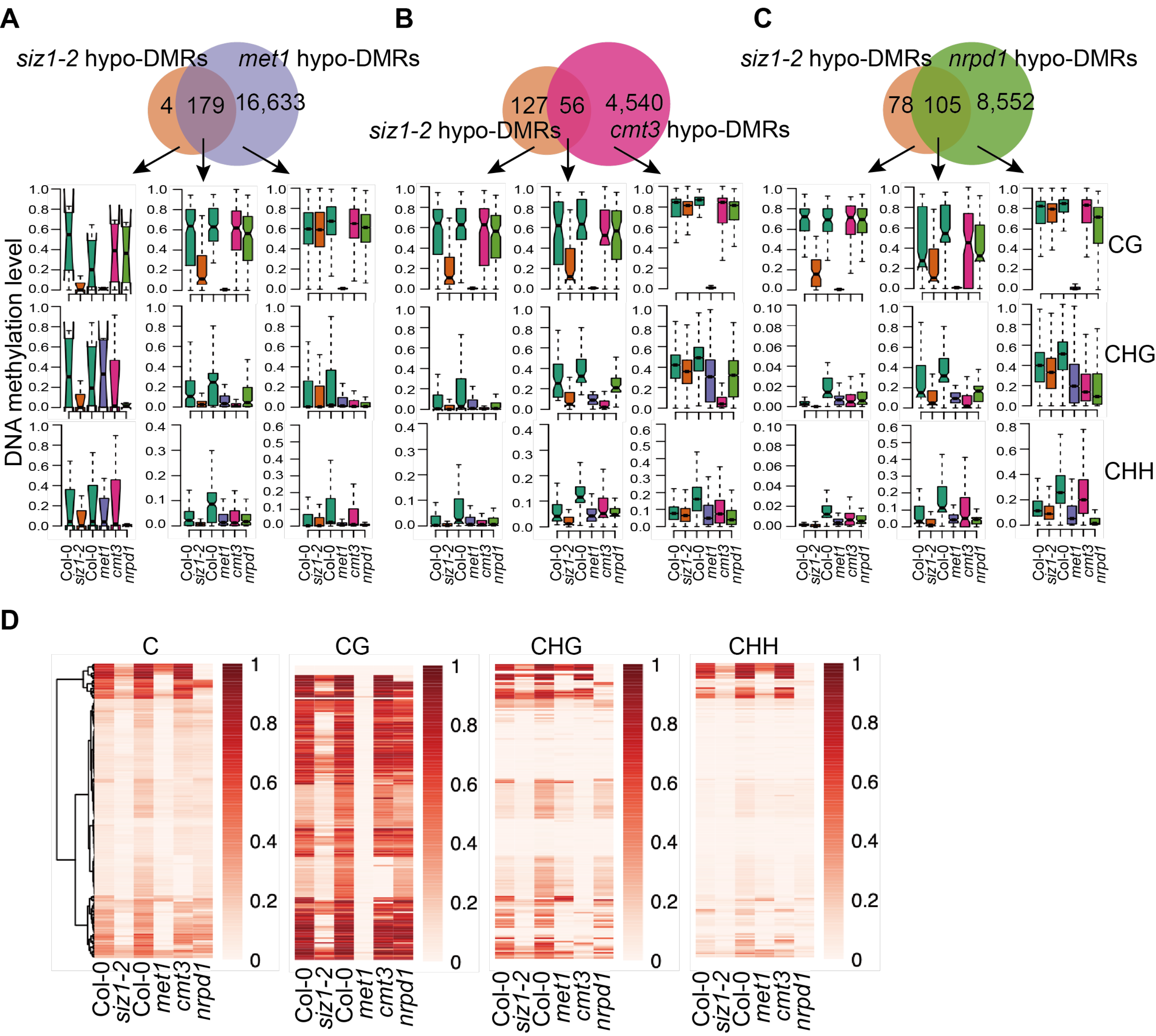


**Supplemental Figure 7. Features of *siz1-2* hypo-DMRs.** (A-C) Venn diagram revealing the numbers of hypo-DMRs that overlapping and unique in *siz1-2* and *met1, cmt3, nrpd1,* and box plots showing the DNA methylation levels at three cytosine sequence contexts that were calculated from the indicated overlapping or unique hypo-DMRs. (D) Heatmap showing the DNA methylation level at different cytosine contexts in *met1, cmt3* and *nrpd1* within those regions that are hypo-methylated in *siz1-2*.

**A**

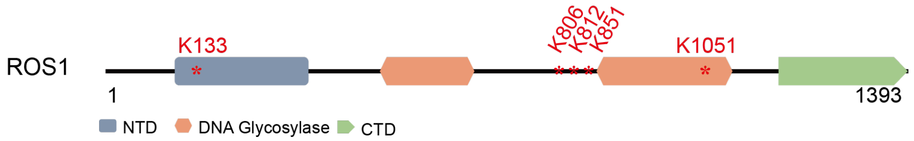


**B**

1 MEKQRREESSFQQPPWIPQTPMKPFSPICPYTVEDQYHSSQLEERRFVGN
51 KDMSGLDHLSFGDLLALANTASLIFSGQTPIPTRNTEVMQKGTEEVESLS
101 SVSNNVAEQILKTPEKPKRKKHRPKVRREAKP**K**REPKPRAPRKSVVTDGQ
151 ESKTPKRKYVRKKVEVSKDQDATPVESSAAVETSTRPKRLCRRVLDFEAE
201 NGENQTNGDIREAGEMESALQEKQLDSGNQELKDCLLSAPSTPKRKRSQG
251 KRKGVQPKKNGSNLEEVDISMAQAAKRRQGPTCCDMNLSGIQYDEQCDYQ
301 KMHWLYSPNLQQGGMRYDAICSKVFSGQQHNYVSAFHATCYSSTSQLSAN
351 RVLTVEERREGIFQGRQESELNVLSDKIDTPIKKKTTGHARFRNLSSMNK
401 LVEVPEHLTSGYCSKPQQNNKILVDTRVTVSKKKPTKSEKSQTKQKNLLP
451 NLCRFPPSFTGLSPDELWKRRNSIETISELLRLLDINREHSETALVPYTM
501 NSQIVLFGGGAGAIVPVTPVKKPRPRPKVDLDDETDRVWKLLLENINSEG
551 VDGSDEQKAKWWEEERNVFRGRADSFIARMHLVQGDRRFTPWKGSVVDSV
601 VGVFLTQNVSDHLSSSAFMSLASQFPVPFVPSSNFDAGTSSMPSIQITYL
651 DSEETMSSPPDHNHSSVTLKNTQPDEEKDYVPSNETSRSSSEIAISAHES
701 VDKTTDSKEYVDSDRKGSSVEVDKTDEKCRVLNLFPSEDSALTCQHSMVS
751 DAPQNTERAGSSSEIDLEGEYRTSFMKLLQGVQVSLEDSNQVSPNMSPGD
801 CSSEI**K**GFQSM**K**EPTKSSVDSSEPGCCSQQDGDVLSCQKPTLKEKGKKVL
851 **K**EEKKAFDWDCLRREAQARAGIREKTRSTMDTVDWKAIRAADVKEVAETI
901 KSRGMNHKLAERIQGFLDRLVNDHGSIDLEWLRDVPPDKAKEYLLSFNGL
951 GLKSVECVRLLTLHHLAFPVDTNVGRIAVRLGWVPLQPLPESLQLHLLEM
1001 YPMLESIQKYLWPRLCKLDQKTLYELHYQMITFGKVFCTKSKPNCNACPM
1051 **K**GECRHFASAFASARLALPSTEKGMGTPDKNPLPLHLPEPFQREQGSEVV
1101 QHSEPAKKVTCCEPIIEEPASPEPETAEVSIADIEEAFFEDPEEIPTIRL
1151 NMDAFTSNLKKIMEHNKELQDGNMSSALVALTAETASLPMPKLKNISQLR
1201 TEHRVYELPDEHPLLAQLEKREPDDPCSYLLAIWTPGETADSIQPSVSTC
1251 IFQANGMLCDEETCFSCNSIKETRSQIVRGTILIPCRTAMRGSFPLNGTY
1301 FQVNEVFADHASSLNPINVPRELIWELPRRTVYFGTSVPTIFKGLSTEKI
1351 QACFWKGYVCVRGFDRKTRGPKPLIARLHFPASKLKGQQANLA

**Supplemental Figure 8. Predicted SUMOylation sites in ROS1.** (A) ROS1 contains a N-terminal domain, a central DNA glycosylase domain with an atypical insertion of approximate 230 amino acids and a C-terminal domain. Predicted SUMOylation sites K133/K806/K812/K851/K1051 are indicated by * in red. By SUMOplot (http://www.abgent.com/sumoplot) and GPS-SUMO (http://sumosp.biocuckoo.org). (B) Amino acids sequence of ROS1. Predicted SUMOylation site lysins (K) are represented in bold and highlighted in red.

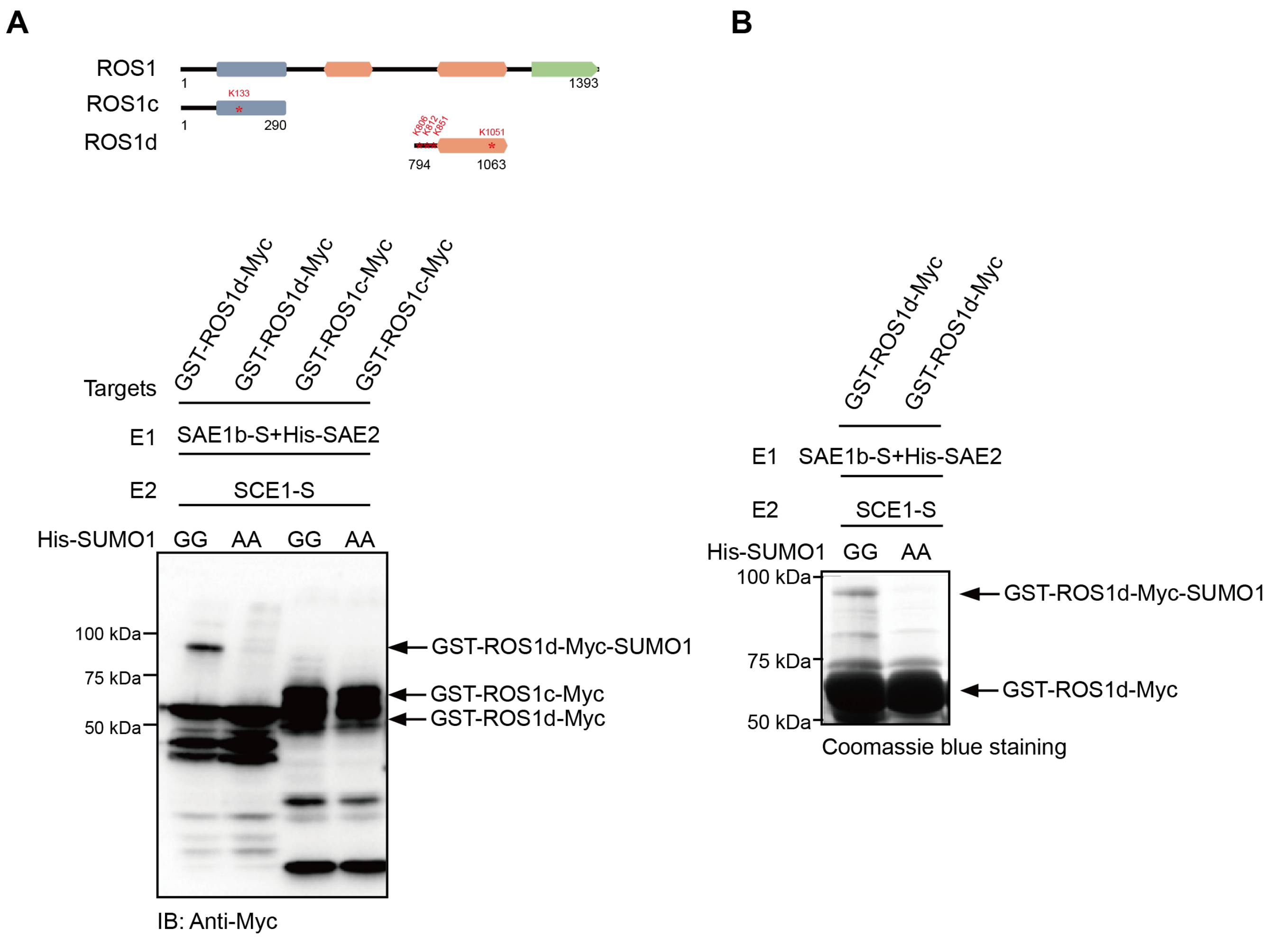


**Supplemental Figure 9. ROS1 is SUMOylated.** (A) SUMOylation of ROS1 was checked via the reconstitution system of Arabidopsis SUMOylation in *E.coli* using Myc-tagged ROS1. Experiment was performed as described in Fig. 4(B). Immuno-blotting was carried out to detect truncated ROS1 and its SUMOylated forms using anti-Myc antibody in crude lysate proteins. Predicted SUMOylation sites K133/K806/K812/K851/K1051 are indicated by * in red. (B) Detection of SUMOylated ROS1 by GST purification. The Coomassie blue  staining of GST purified proteins was shown.

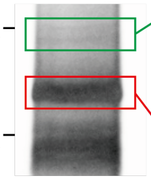


**ROS1 Purification (Anti-Flag)**

| Proteins | Accession number | Unique peptide number | Coverage / % |
|---|---|---|---|
| ROS1 | At2g36490 | 48 | 38.5 |
| SUMO1 | At4g26840 | 1 | 10.0 |

| Proteins | Accession number | Unique peptide number | Coverage / % |
|---|---|---|---|
| ROS1 | At2g36490 | 72 | 53.0 |
| SUMO1 | At4g26840 | N.D. | N.D. |

**Supplemental Figure 10. Detection of SUMOylated ROS1 by LC-MS/MS.** LC-MS/MS following immunoprecipitation using an anti-Flag antibody for immuno-purifying ROS1-Flag co-expressed with HA-SUMO1 in Col-0 protoplast. The immuno-purified proteins was separated by SDS-PAGE and silver stained (left panel), the ROS1-Flag (in red rectangle) and SUMOylated ROS1-Flag (in green rectangle) bands were cut for LC-MS/MS. The selected proteins were listed in the right panel. N.D., not detected.

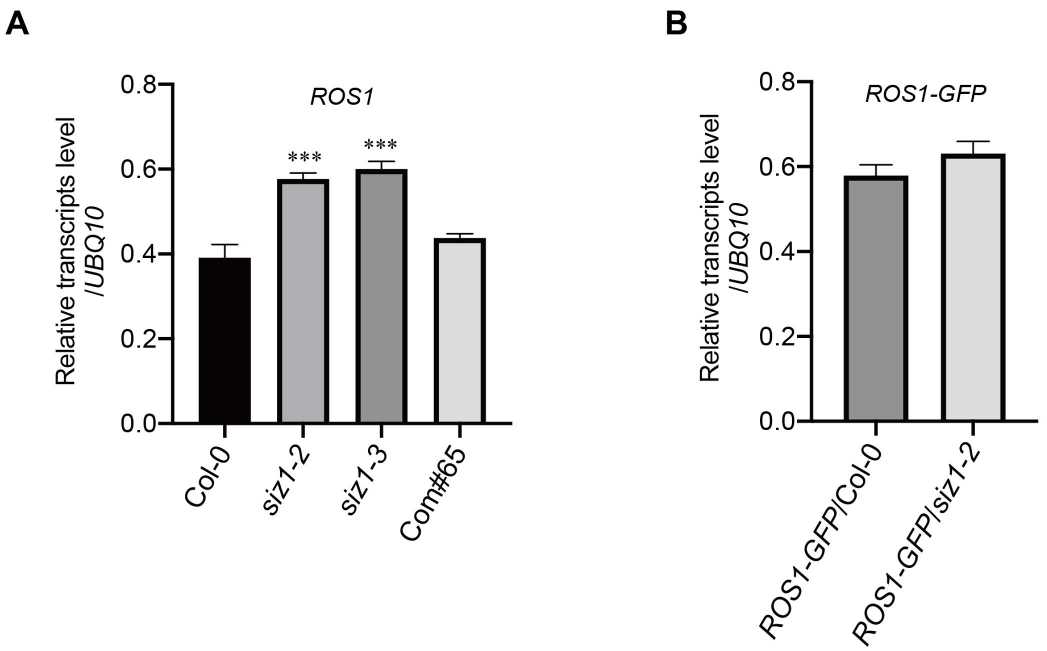


**Supplemental Figure 11. *ROS1* transcript level in the *siz1* mutants.** (A) Relative transcripts level of *ROS1* in *siz1* mutants was determined by RT-qPCR. Compared to Col-0, the transcripts level of *ROS1* in *siz1* mutants was significant increased, which was rescued by complementation of *SIZ1* in *siz1-2*. The error bars indicate the mean ± SD (n=3). The asterisks show significant differences between Col-0 and indicated mutants, $^{***}P < 0.001$, Student's *t*-test. (B) Relative transcripts level of *ROS1-GFP* in knock-in lines was determined by RT-qPCR. The *ROS1-GFP* transcripts level was relatively higher in *siz1-2*. The error bars indicates the mean ± SD (n=3).

**Supplemental Table 3.** Primers used in this study.

| **Primers for genotyping and RT-qPCR** | |
|---|---|
| **Primer Name** | **Primers Sequences** |
| *siz1*-LP | GAGCTGAAGCATCTGGTTTTG |
| *siz1*-RP | CACGACAGATGAAGCATTGTG |
| LBb1.3 | ATTTTGCCGATTTCGGAAC |
| *SIZ1*-qRT-F | GCTGACGTTTCAGGAGGTTTAGTTG |
| *SIZ1*-qRT-R | GCCTTGTCTTGTCTACTGTCATTCATAC |
| *TUB8*-qRT-F | ATAACCGTTTCAAATTCTCTCTCTC |
| *TUB8*-qRT-R | TGCAAATCGTTCTCTCCTTG |
| *UBQ10*-qRT-F | GATCTTTGCCGGAAAACAATTGGAGGATGGT |
| *UBQ10*-qRT-R | CGACTTGTCATTAGAAAGAAAGAGATAACAGG |
| *ACT2*-qRT-F | TGCCAATCTACGAGGGTTTC |
| *ACT2*-qRT-R | TTACAATTTCCCGCTCTGCT |
| *ROS1*-qRT-F | AAGGTCACATGTTGTGAACCAAT |
| *ROS1*-qRT-R | ATGCTCGTCTGGAAGTTCGTA |
| *At1g28480*-qRT-F | ACGGAGAGGATGTTGCATGTGTC |
| *At1g28480*-qRT-R | AATCTCAAGGACCGCCGGATTC |
| *At1g58270*-qRT-F | CGTTTTCTTGACTCTTACACTTCTGA |
| *At1g58270*-qRT-R | CCATTTGGATACACTTTCAATGC |
| *At2g30810*-qRT-F | CAAGCTCAAGCACCTGCAATCC |
| *At2g30810*-qRT-R | ACATGCCTTTGGACATTCTTCTGG |
| *At3g27480*-qRT-F | TGACTCGCCAACAGCTTGTTCTG |
| *At3g27480*-qRT-R | GGCACATCAATGGAACTGCATAGC |
| *At4g21030*-qRT-F | AGTGTGTCCAAGGTGTTATTCCG |
| *At4g21030*-qRT-R | GCGCGGTTGAGACTTCTTGTTG |
| *At5g45930*-qRT-F | TGTGCTGAGCTGGACGTTGATG |
| *At5g45930*-qRT-R | AGCGCTCTAGCTGCTCTGTTTATC |

| **Primers for CHOP-PCR and locus-specific bisulfite sequencing** | |
|---|---|
| **Primer Name** | **Primers Sequences** |
| *At1g26390*-CHOP-F | GAACCTGAGTGCTGAAGGTTACTC |
| *At1g26390*-CHOP-R | CAAGATTCAATACTTTACGACATG |
| *At1g26400*-CHOP-F | TGACCTGCATAGGCTATAACACA |
| *At1g26400*-CHOP-R | ATTGGAATCAATCCGAGTGG |
| *At1g26410*-CHOP-F | GTACGTTTACACGTGTATTTTATT |
| *At1g26410*-CHOP-R | GTAGCGTCTCGTTGCAATTCAAC |
| *At4g18650*-CHOP-F | TCCACATGAGCCATCAACTC |
| *At4g18650*-CHOP-R | AAGGGTACCAGTTTGGGAAAA |
| HDM1-CHOP-F | GCATACACATCATGCGAC |
| HDM1-CHOP-R | GAAGCAACAATTATGGTAC |
| HDM2-CHOP-F | GATTAATCATGTACCCTTC |

| | |
|---|---|
| HDM2-CHOP-R | GGAGAATTGTCTGATTTGTAAC |
| HDM3-CHOP-F | ACTTGCAAATGAAGACTTAG |
| HDM3-CHOP-R | CTTGGTTCTTAAACATTTG |
| HDM4-CHOP-F | GGATTCGTGGATCCAAGTCA |
| HDM4-CHOP-R | GCATTATATTCGATGTGATG |
| HDM5-CHOP-F | CTCAACAAGGTCAACATGTAGC |
| HDM5-CHOP-R | CAGGATTCATGCACTTTCC |
| HDM6-CHOP-F | TGCTCCACTTAGAAGTCCT |
| HDM6-CHOP-R | TTGGTATCGTCGACGGAG |
| HDM7-CHOP-F | GAACCTCCAAACTGAACAC |
| HDM7-CHOP-R | GCCAAATATCTTACTACACC |
| HDM8-CHOP-F | GGATTCATCTACAACGACGT |
| HDM8-CHOP-R | GGAGTTAATGTAATCGTGAG |
| HDM9-CHOP-F | GCCTATCTACCAACTTGAC |
| HDM9-CHOP-R | GTATTCAATTGCTACGTGGA |
| *At1g26390*-bis-F1 | TTAGGGTTAAAATATGAAGATTGTTAAGAAATGAGTTGG |
| *At1g26390*-bis-R1 | ATCATACTTAACTTTAACATCCATCAATCTCTTC |
| *At1g26390*-bis-F2 | GAAGATTGTTAAGAAATGAGTTGGTTTAATTTAAYG |
| *At1g26390*-bis-R2 | TCAATCTCTTCAAATTCCCCAAAAAATACTT |
| *At1g26400*-bis-F1 | TATTTTGTCGATTTCATTTGTTGG |
| *At1g26400*-bis-R1 | CCACAACATTTCCTCACC |
| *At1g26400*-bis-F2 | GTAGTTTGAGATGATTAATGATAGAGTT |
| *At1g26400*-bis-R2 | AAACTTATTCAATCTTCAATACTCTAC |
| *At1g26410*-bis-F1 | CACATACCCAAATAACTTAATATCACTACTTTACACR |
| *At1g26410*-bis-R1 | GAGAAGTTTTGGAAGATAATGTTTAAATTTAATAGTA |
| *At1g26410*-bis-F2 | CCCAAATAACTTAATATCACTACTTTACACAAACC |
| *At1g26410*-bis-R2 | GGAAGATAATGTTTAAATTTAATAGTAGTGT |

### Primers for cloning

| **Primer Name** | **Primers Sequences** |
|---|---|
| SIZ1-AD-F | CAGATTACGCTCATATGATGGATTTGGAAGCTAATTG |
| SIZ1-AD-R | TCGAGCTCGATGGATCCTTAAACTCCGGTGTCTTGTC |
| ROS1-BD-F | AGGAGGACCTGCATATGATGGAGAAACAGAGGAGAG |
| ROS1-BD-R | TGCAGGTCGACGGATCCTTAGGCGAGGTTAGCTTGTTG |
| ROS1-nLUC-F | CGGGGGACGAGCTCGGTACCATGGAGAAACAGAGGAGAGAAGAAAGC |
| ROS1-nLUC-R | GCGTACGAGATCTGGTCGACGGCGAGGTTAGCTTGTTGTCCCTT |
| ROS1a-nLUC-F | CGGGGGACGAGCTCGGTACCATGCCAAAGAAGAAGCGGAAGGTCATGGAGAAACAGAGGAGAGAAG |
| ROS1a-nLUC-R | GCGTACGAGATCTGGTCGACACCACCAAAGAGTACAATC |
| ROS1b-nLUC-F | CGGGGGACGAGCTCGGTACCATGCCAAAGAAGAAGCGGAAGGTCGGCGCTGGAGCAATTGTG |
| ROS1b-nLUC-R | GCGTACGAGATCTGGTCGACGGCGAGGTTAGCTTGTTGTC |
| cLUC-SIZ1-F | ACGCGTCCCGGGGCGGTACCATGGATTTGGAAGCTAATTG |

| | |
|---|---|
| cLUC-SIZ1-R | CGAAAGCTCTGCAGGTCGACTTAAACTCCGGTGTCTTGTC |
| cLUC-SUMO1-F | ACGCGTCCCGGGGCGGTACCATGTCTGCAAACCAGGAGG |
| cLUC-SUMO1-R | CGAAAGCTCTGCAGGTCGACTCAGGCCGTAGCACCACCAC |
| ROS1-nYFP-F | CGAACGATAGTTAATTAAATGGAGAAACAGAGGAGAG |
| ROS1-nYFP-R | GCCACCTCCTCCACTAGTGGCGAGGTTAGCTTGTTGTCCC |
| SIZ1-cYFP-F | CGAACGATAGTTAATTAAATGGATTTGGAAGCTAATTGTAAGG |
| SIZ1-cYFP-R | GCCACCTCCTCCACTAGTAACTCCGGTGTCTTGTC |
| SIZ1-pUC18HA-F | CATCCTCAATTTGAAAAAGGATCCATGGATTTGGAAGCTAATTGTAAGG |
| SIZ1-pUC18HA-R | GAGCTCTATCGATCAATCATTAAACTCCGGTGTCTTGTC |
| ROS1-pUC18Flag-F | GATGACGATAAGGGATCCATGGAGAAACAGAGGAGAG |
| ROS1-pUC18Flag-R | ATCGATCAATCAGGATCCTTAGGCGAGGTTAGCTTGTTGTC |
| ROS1b-pUC18Flag-F | ATTTACGAACGATACTCGAGATGGGCGCTGGAGCAATTGTG |
| ROS1b-pUC18Flag-R | TCTTTGTAGTCCATGTCGACGGCGAGGTTAGCTTGTTGTC |
| ROS1c-pET28a-F | GGTCGCGGATCCGAATTCATGGAGAAACAGAGGAGAGAAG |
| ROS1c-pET28a-R | GTGGTGGTGGTGCTCGAGCCCTGATAGATTCATGTCGC |
| ROS1d-pET28a-F | GGTCGCGGATCCGAATTCCCAAATATGTCTCCGGGTGATTG |
| ROS1d-pET28a-R | GTGGTGGTGGTGCTCGAGACTTGCAAACGCACTGGC |
| ROS1cMyc-pGEX4T1-F | TCTACGAATTCATGGAGAAACAGAGGAGAGAAG |
| ROS1cMyc-pGEX4T1-R | TCTACCTCGAGTTATTCATTCAAGTCCTCTTCAGAAATGAGCTTTTGCTCCATCCCTGATAGATTCATGTCGC |
| ROS1dMyc-pGEX4T1-F | TCTACGAATTCATGCCAAATATGTCTCCGGGTGATTG |
| ROS1dMyc-pGEX4T1-R | TCTACCTCGAGTTATTCATTCAAGTCCTCTTCAGAAATGAGCTTTTGCTCCATACTTGCAAACGCACTGGC |
| SCE-1305Flag-F | TCTACGGTACCATGGCTAGTGGAATCGCTCGT |
| SCE-1305Flag-R | TCTACAAGCTTGACAAGAGCAGGATACTGCTTGGA |

### Primers for site direct mutagenesis

| **Primer Name** | **Primers Sequences** |
|---|---|
| ROS1-K133R-F | GCTAAACCCAGGAGGGAGCCTAAACCACGAG |
| ROS1-K133R-R | AGGCTCCCTCCTGGGTTTAGCTTCTCTACGA |
| ROS1-K806R-F | TCAGAAATTAGGGGTTTCCAGTCAATGAAAG |
| ROS1-K806R-R | CTGGAAACCCCTAATTTCTGAGCTACAATCA |
| ROS1-K812R-F | CAGTCAATGAGAGAGCCCACAAAATCCTCTG |
| ROS1-K812R-R | TGTGGGCTCTCTCATTGACTGGAAACCCTTA |
| ROS1-K851R-F | AAGGTTTTGAGGGAGGAAAAAAAAGCGTTTG |
| ROS1-K851R-R | TTTTTCCTCCCTCAAAACCTTTTTCCCTTTT |
| ROS1-K1051R-F | TGTCCGATGAGAGGAGAATGCAGACATTTTG |
| ROS1-K1051R-R | GCATTCTCCTCTCATCGGACATGCATTGCAA |
| SCE-C94S-F | TGGAACTGTCAGTCTCTCTATCCTTAACGAGGATTATGGATGGAGACC |
| SCE-C94S-F | GGATAGAGAGACTGACAGTTCCAGATGGATAGACATTAGGGTGGAA |
